# Trends and projections of universal health coverage indicators in Ghana, 1995-2030: A national and subnational study

**DOI:** 10.1101/486159

**Authors:** Cherri Zhang, Md. Shafiur Rahman, Md. Mizanur Rahman, Alfred E Yawson, Kenji Shibuya

## Abstract

Ghana has made significant stride towards universal health coverage (UHC) by implementing the National Health Insurance Scheme (NHIS) in 2003. This paper investigates the progress of UHC indicators in Ghana from 1995 to 2030 and makes future predictions up to 2030 to assess the probability of achieving UHC targets. National representative surveys of Ghana were used to assess health service coverage and financial risk protection. The analysis estimated the coverage of 13 prevention and four treatment service indicators at the national level and across wealth quintiles. In addition, this analysis calculated catastrophic health payments and impoverishment to assess financial hardship and used a Bayesian regression model to estimate trends and future projections as well as the probabilities of achieving UHC targets by 2030. Wealth-based inequalities and regional disparities were also assessed. At the national level, 14 out of the 17 health service indicators are projected to reach the target of 80% coverage by 2030. Across wealth quintiles, inequalities were observed amongst most indicators with richer groups obtaining more coverage than their poorer counterparts. Subnational analysis revealed while all regions will achieve the 80% coverage target with high probabilities for prevention services, the same cannot be applied to treatment services. In 2015, the proportion of households that suffered catastrophic health payments and impoverishment at a threshold of 25% non-food expenditure were 1.9% (95%CrI: 0.9-3.5) and 0.4% (95%CrI: 0.2-0.8), respectively. These are projected to reduce to less than 0.5% by 2030. Inequality measures and subnational assessment revealed that catastrophic expenditure experienced by wealth quintiles and regions are not equal. Significant improvements were seen in both health service coverage and financial risk protection as a result of NHIS. However, inequalities across wealth quintiles and at the subnational level continue to be cause of concerns. Further efforts are needed to narrow these inequality gaps.

## Introduction

Universal Health Coverage (UHC) is a concept in which all people receive the quality, essential services they need without experiencing financial hardship [1,2]. The First Global Monitoring Report formulated by World Health Organizations (WHO) and World Bank identified three dimensions: population, health services, and financing through risk pooling mechanism to track UHC progress [1]. Since its integration into the recently adopted Sustainable Development Goal (SDG) 3, member countries of the United Nations (UN) have committed to achieve UHC by 2030 [3]. This commitment consists of two targets: a minimum of 80% essential health service coverage for all people, regardless of socioeconomic status, and 100% financial risk protection from out-of-pocket (OOP) payments for health care [1]. UHC is a key mechanism to ensure affordability and equity as well as to guarantee resilient health system, which many countries have embraced in order to achieve better health for all [1].

Achieving UHC in Sub-Saharan Africa should be of utmost priority as countries in this region trail significantly behind in achieving health outcomes especially the Millennium Development Goals (MDGs) formulated by the WHO. Moreover, millions of Africans fall into poverty annually due to OOP payment as a result of lack of health insurance in health financing system [4,5]. Ghana, being one of the few Sub-Saharan African countries advocating for UHC, implemented the National Health Insurance Scheme (NHIS) in 2003, in an attempt to remove financial barriers, protect Ghanaians from catastrophic expenditure, and improve access for everyone [6]. NHIS is mainly funded by three sources: 70% of National Insurance Levy (NIL), 17.4% of Social Security and National Insurance Trust (SSNIT), and 4.5% of premium payments [7]. NIL is a 2.5% tax on selected goods and services; SSNIT is a 2.5% contribution paid by those in the formal sectors, and premium is set at an annual flat rate of $4.8 USD to $32 USD depending on districts for those in the informal sectors [8]. Pregnant women and those under the age of 18 years or over 70 years of age are exempt from premium and makes up 60% of the enrollees [8].

Services covered under the insurance include outpatient and inpatient care, oral health, eye care, maternity care, and emergencies with no copayment upon receipt of service. It excludes cosmetic services, HIV antiretroviral drugs, orthopedics, and organ transplant etc. [9]. Despite enrollment into NHIS being mandatory, overall enrollment remains low at 40% as of 2016 since majority of the population belongs to the informal sector, and there is a lack of formal tracking regulations [7]. Although Ghana made significant stride towards health financing in recent years; however, Gross Domestic Product (GDP) spending on health and total government expenditure allocated to health has dropped since 2010 to 3.6% and 6.8%, respectively in 2014 [10]. Furthermore, other challenges such as funding and sustainability persist as overall enrollment has decreased in recent years and many citizens find paying for premiums difficult. In addition, as the incidence of poverty is around 25% with some regions such as Upper East and Upper West experiencing more than 70% incidence of poverty, inequality remains a prominent issue especially across regions [9]. Inadequate funding for health can lead to unstable health insurance scheme and an increase in OOP payments pushing people further into poverty and resulting in worse health outcomes.

Most studies conducted in Ghana thus far assessed the impact of NHIS either on health service access or protection against financial catastrophe; therefore, this paper offers a comprehensive glimpse into Ghana’s progress towards both UHC components, along with future trajectories. It will provide a thorough examination on the proportion of health service utilization as well as catastrophic health expenditures at the national and subnational level as well as across wealth quintiles using nationally representative survey data.

## Methods

### Data source

Analysis on health service coverage was carried out using five consecutive Demographic and Health Surveys (1993, 1998, 2003, 2008, and 2014). These are nationally representative household surveys covering all 10 regions of Ghana. Information mainly on housing and household characteristics, education, maternal and child health, fertility, family planning, and nutrition were collected The analysis of financial risk protection was conducted using Ghana Living Standard Surveys (1991-1992, 1998-1999, 2005-2006, and 2012-2013), which are nationwide household surveys constructed to gather information on living conditions in Ghana. The surveys collected detailed information on household demographics, education, health, employment, migration, housing conditions, agriculture, and access to financial services and asset ownership. Both national surveys involved two-staged random sampling design with high response rates (Table in S11 Table).

### Measurement of health service indicators

In accordance with WHO and World Bank’s framework [1], 17 health service indicators (Table in S1 Table) were chosen to cover a full range of prevention and treatment services based on data availability. In all, 13 indicators were classified as prevention services while four indicators were classified as treatment services. The chosen indicators were grouped into: 1) composite prevention index and 2) composite treatment index. These were estimated as the mean value of prevention and treatment service indicators to trace the overall progress of prevention and treatment coverage [11]. Due to incomplete data for some of the indicators in certain survey years, only nine out of the 13 prevention indicators (four antenatal care visits, exclusive breastfeeding, needs for family planning satisfaction, improved water, adequate sanitation, BCG, measles, DPT3 and Polio3 immunizations) were included to estimate the composite prevention index. Furthermore, a Composite Coverage Index (CCI) was estimated to assess access to maternal and child health services as it frequently represents frontline measurement of health service coverage and produce the most immediate picture of accessibility [11,12]. It was calculated based on eight interventions from four specialties (family planning, maternity care, child malnutrition, and case management) using a formula developed by Boerma and colleagues [12].

### Measurement of financial hardship

Incidence of catastrophic health expenditure (CHE) and impoverishment due to OOP health payments were approximated for financial hardship assessment. Several thresholds can be considered according to World Bank’s guideline [13]. In this analysis, a threshold of 25% non-food consumption expenditure was used; therefore, a household’s health expenditure was deemed catastrophic if its total OOP health payments exceeded that threshold. Consistent with WHO’s guideline [14], the incidence of impoverishment was derived using poverty line and subsistence spending. A household was considered poor if its total per capita expenditure was less than its subsistence spending after paying for healthcare, and thus a household’s health expenditure was considered impoverishing if its total per capita spending after paying for health care was below the poverty line [14]. The poverty line was determined using average food expenditure of households with food expenditure share within 45th and 55th percentile of the sampled households [14]. A household was deemed to be experiencing hardship if it encountered either CHE or impoverishment. All household consumption calculations were performed following Living Standard Measurement Study guideline [13].

### Statistical analysis

All trends and projections for health service coverage and financial risk protection components were estimated based on proportions estimated from original survey data with 95% confidence interval (CI). The composite prevention and treatment indices were developed based on random-effects meta-analysis [15]. For equity analysis, households were divided into five wealth quintiles (Q1-Q5) to assess socio-economic status. Due to the lack of information on income as most Ghanaians belong to the informal sector, household economic status was measured based on the level of consumption or asset-based wealth index. The index was constructed from household asset data using principle components analysis and a wealth score was generated for each household. This information was provided for households in the Demographic and Health Surveys as well as Ghana Living Standard Measurement Surveys. Households were ranked based on wealth scores and divided into quintiles, starting from the poorest quintile (lowest 20%) to the richest quintile (highest 20%). The slope index of inequality (SII) and relative index of inequality (RII) were calculated to provide an absolute and relative measure of inequality, respectively. SII measures the absolute difference between the extremes of wealth quintiles and reflects the difference in percentage points in each indicator, while RII is a measure of ratio signifying the degree of inequality [11]. SII and RII were calculated by regressing outcomes of health service and financial indicators against household’s relative rank in the cumulative distribution of wealth position. All aforementioned analyses were performed using Stata (version 15.0/MP, StataCorp).

A Bayesian linear regression model with a non-informative prior was developed, considering year as the covariate, to estimate the trends in indicators over time and its posterior predictive distribution. All proportions were logit transformed before the analysis, and all calculations were conducted as such. The Markov Chain Monte Carlo (MCMC) algorithm was applied to obtain 1000 samples from the posterior distribution of the parameter of interest using two chains. For each of the chain, the first 5000 iterations were discarded as burn-ins and the number of iterations increased until the MCMC outputs converged. These posterior predictive distributions were used to obtain projections and credible intervals up to year 2030. They were also utilized to calculate the annual rate of change and the probability of achieving UHC targets for all included indicators. Another wealth quintile adjusted model with non-informative prior was fitted to estimate the predicted coverage of all indicators for different socio-economic groups.

Convergence of MCMC outputs was assessed by visually examining trace plots. Posterior samples were considered to have converged when outputs from two chains adjoined.

Additionally, Gelman-Rubin diagnostic statistics were applied as a quantifiable measure of convergence. A potential scale reduction factor (PSRF) was used in the Gelman diagnostic, where a PSRF value close to 1 signified convergence, and a PSRF value greater than 1.02 indicated convergence failure. To further assess the accuracy of the model, a deviance information criterion (DIC) was calculated for each indicator at the wealth quintile and subnational level in the case that quintiles or regions also acted as covariates. For every single estimate of trend and projection in health service coverage, a DIC value was calculated for model with and without interaction. The model with smaller penalized deviance was integrated into the analysis (Table in S2 Table). Bayesian regression models were developed in JAGS and implemented in R.

## Results

### Health service coverage

Table 1 lists the predicted coverage of 17 chosen health service indicators, grouped into prevention and treatment categories along with 95% credible intervals (CrI), the probability of achieving the target of 80% coverage by the year 2030, as well as the annual rate of change from 1995 to 2030. The results are very promising amongst most prevention service indicators as they are estimated to have more than 90% probability of achieving the target except need for family planning satisfied, adequate sanitation, and non-use of tobacco. Amongst the treatment indicators, care seeking for pneumonia among children is the least reassuring with a low probability of 5.1% of reaching the target while access to institutional delivery and use of skilled birth attendance will have more than 85% probability. In 2015, family planning demand satisfied and care seeking for pneumonia were deemed to have the lowest national coverage at 46.3% (95% CrI: 38.5-54.3) and 49.9% (95% CrI: 33.8-64.0), respectively, followed by insecticide treated bed nets for pregnant women and children at around 60%.

**Table 1:** National health service coverage with probability of achieving the target and rate of change, 1995-2030.

| Indicators | Predicted coverage in year (95% CrI) |  |  |  | Probabilit<br>y<sup>a</sup> | Annual % change<sup>b</sup> |
| --- | --- | --- | --- | --- | --- | --- |
|  | 1995 | 2005 | 2015 | 2030 |  |  |
| Prevention indicators |  |  |  |  |  |  |
| Needs for family<br>planning satisfied | 36.6 (30.5-42.7) | 41.4 (37.2-45.7) | 46.3 (38.5-54.3) | 53.7 (38.4-68.9) | 0.9% | 1.1 (-0.2-2.3) |
| At least four antenatal<br>care visits | 57.9 (49.6-63.5) | 73.7 (69.3-77.8) | 84.9 (78.3-89.4) | 93.9 (87.5-97.5) | 99.9% | 1.4 (0.9-2.0) |
| Postnatal care for<br>mothers | 0.4 (0.1-1.3) | 10.6 (5.6-17.6) | 79.9 (61.7-91.3) | 99.8 (99.0-100.0) | 100% | 18.6 (13.5-23.9) |
| Exclusive breastfeeding | 16.6 (9.6-25.2) | 40.7 (31.2-50.5) | 70.1 (52.3-83.2) | 92.6 (77.1-98.5) | 96.3% | 5.3 (3.4-7.1) |
| Insecticide treated bed<br>nets for children | – | 10.5 (5.9-17.0) | 62.7 (42.7-80.4) | 98.1 (91.1-99.9) | 99.5% | 10.4 (7.5-13.2) |
| Insecticide treated bed<br>nets for mothers | – | 6.7 (3.8-10.7) | 56.7 (37.2-74.4) | 98.3 (91.8-99.9) | 99.6% | 12.2 (9.2-15.1) |
| BCG immunization | 84.9 (80.4-88.6) | 93.4 (92.1-94.7) | 97.2 (95.9-98.2) | 99.2 (98.4-99.7) | 100% | 0.5 (0.3-0.7) |
| DPT3 immunization | 67.4 (59.9-75.0) | 82.1 (78.0-85.6) | 90.9 (86.3-94.3) | 96.8 (93.2-98.8) | 100% | 1.1 (0.6-1.5) |
| Polio3 immunization | 67.9 (58.9-76.2) | 79.6 (75.1-83.9) | 87.6 (81.8-92.4) | 94.2 (87.3-98.0) | 100% | 1.0 (0.5-1.5) |
| Measles immunization | 69.5 (60.6-77.9) | 83.8 (79.9-87.5) | 92.0 (87.5-95.1) | 97.3 (93.4-99.1) | 100% | 1.0 (0.6-1.5) |
| Improved water | 60.9 (50.9-70.4) | 77.0 (71.7-82.0) | 87.6 (80.9-92.6) | 95.3 (88.7-98.5) | 100% | 1.4 (0.7-2.0) |
| Adequate Sanitation | 24.6 (18.7-32.8) | 43.2 (37.8-49.4) | 64.1 (53.3-73.7) | 86.0 (72.4-94.1) | 88.4% | 3.8 (2.6-4.8) |
| No-use of tobacco | – | 91.5 (87.6-94.4) | 91.1 (85.3-95.2) | 88.8 (62.9-98.3) | 87.5% | -0.1 (-1.0-0.6) |
| **Treatment indicators** |  |  |  |  |  |  |
| Institutional delivery | 40.3 (31.2-50.8) | 56.7 (49.6-63.2) | 71.5 (59.3-81.6) | 86.3 (69.9-95.1) | 86.2% | 2.3 (1.1-3.3) |
| Skilled birth attendance | 41.3 (31.6-52.0) | 57.5 (50.4-63.8) | 72.1 (61.0-81.4) | 86.6 (70.7-95.2) | 88.1% | 2.2 (1.2-3.2) |
| Oral rehydration therapy | 39.5 (27.8-51.6) | 55.6 (46.9-64.0) | 70.5 (55.8-82.1) | 85.2 (63.4-95.9) | 79.5% | 2.3 (0.8-3.5) |
| Care seeking for pneumonia | 35.9 (25.9-46.6) | 42.7 (34.6-50.2) | 49.9 (33.8-64.0) | 59.9 (30.1-83.6) | 5.1% | 1.5 (-0.6-3.3) |

Note: <sup>a</sup>The probability of meeting the target of 80% health service coverage by 2030 for the entire population, regardless of economic status, gender, or place of residence according to WHO's universal health coverage target. <sup>d</sup> The annual rate of change for the period 1995-2030. Trends and projections for all the year from 1993-2030 at the national level and across wealth quintiles for all indicators are shown in appendix (p13-21). CrI= credible interval; DPT3= three doses of DPT immunization; Polio3= three doses of polio immunization.

A notable achievement is that national coverage of the four childhood vaccinations (BCG, measles, three doses of DPT and polio vaccination) had already reached the target in 2015. In addition, coverage for maternal postnatal care increased from 10.6% (95%CrI: 5.6-17.6) in 2005 to 79.9% (95% CrI: 61.7-91.3) in 2015. The lowest coverage among the poorest quintile was observed in adequate sanitation at 6.5% (95% CrI:3.4-11.1) in 2015. This is followed by need for family planning demand satisfied at 38.1% (95%CrI: 31.1-45.1) and access to skilled birth attendance at 38.9% (95% CrI: 25.3-53.7). All quintiles will fail to achieve the 80% coverage target for family planning demand satisfied and care seeking for pneumonia. Besides the two aforementioned indicators, the richest quintile is highly likely to reach the target for most indicators except polio vaccination and insecticide treated bed nets use by children under five. A detailed breakdown of coverage by wealth quintile for each indicator can be found in Tables in S3-S6 Table and Figures in S2-S10 Figure.

In order to provide a broader picture regarding the coverage of prevention and treatment services, the composite prevention and treatment indices (Figure 1 and 2) illustrate that overall national coverage of prevention and treatment services are expected to reach 92.2% (95% CrI: 85.4-96.5) and 80.3% (95% CrI: 67.7-89.4) by 2030 and all quintiles other than the two poorest quintiles in treatment index will have high probabilities of achieving the 80% coverage target. Trends and projections in CCI related to reproductive, maternal and child health indicators between 1993 and 2030 states that overall national coverage will increase to 80.7% (95%CrI: 77.3-83.9) by 2030 (Table in S7 Table and Figure in S1 Figure); however, only the middle class and the richer quintile are predicted to achieve the target. Detailed quintile-specific coverage of all three composite indices with probabilities of reaching the 80% coverage target by 2030 is presented in Table is S7 Table.

**Figure 1:**
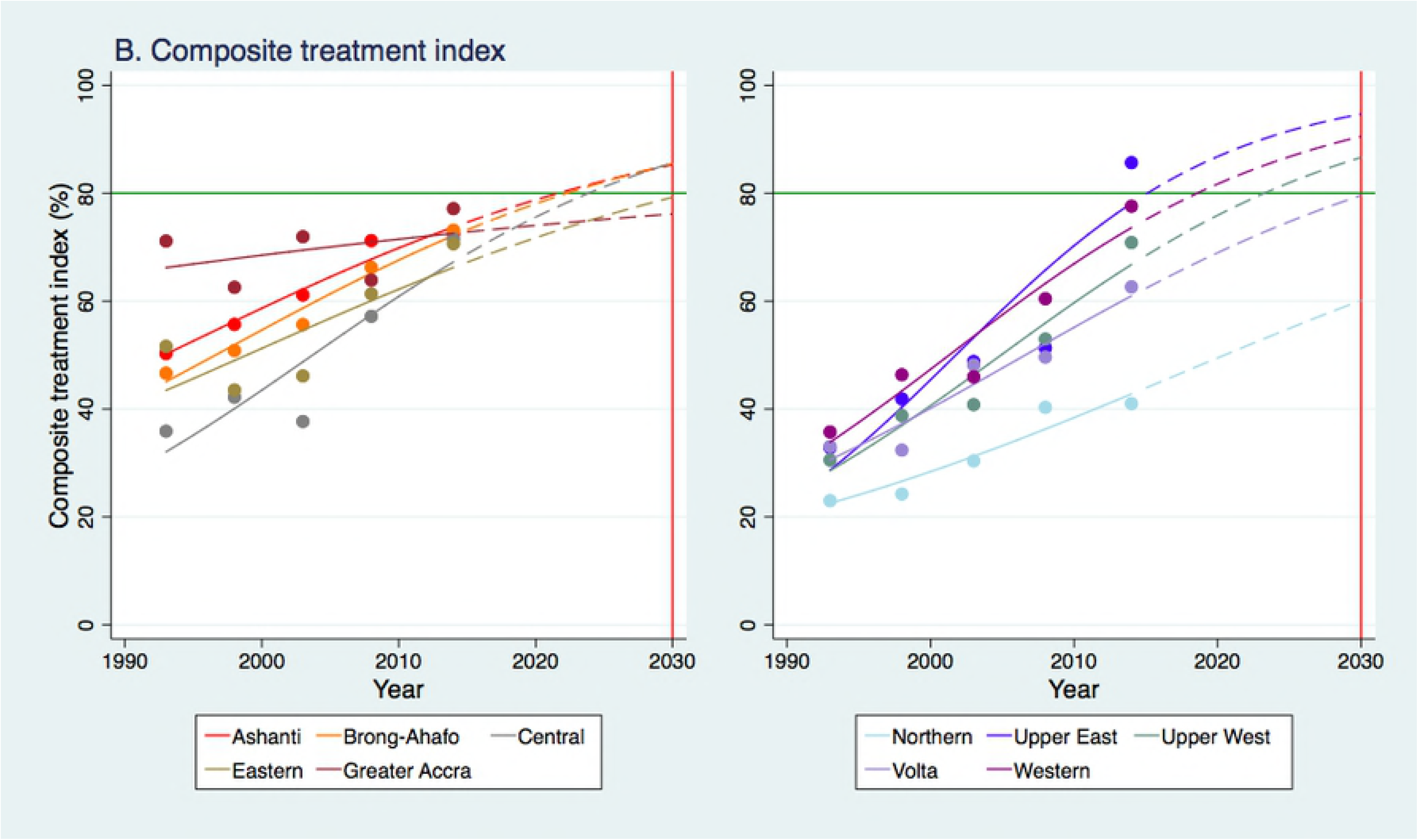
Trends in and projections of overall prevention coverage in Ghana, 1993-2030.

**Figure 2:**
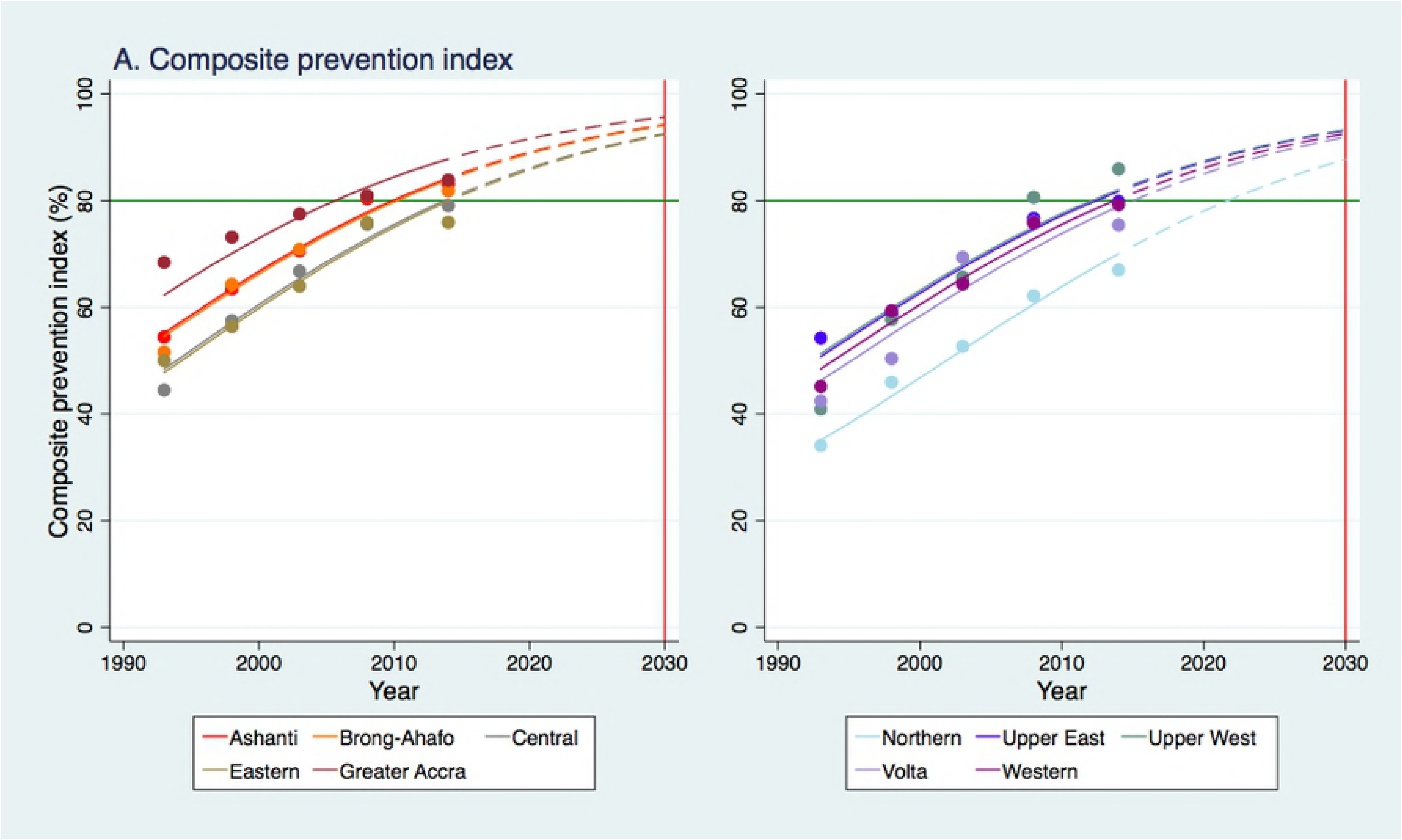
Trends in and projections of overall treatment coverage in Ghana, 1993-2030.

Figure 3 and 4 presents the trends and projections of prevention and treatment indices across the 10 regions in Ghana from 1995 to 2030. All coverage are shown to have increased and are predicted to continue up to 2030. In 2015, the lowest overall prevention and treatment service coverage were observed in Northern at 71.5% (95%CrI: 67.7-75.2) and 43.9% (95%CrI: 33.2-55.1) respectively, while the highest overall coverage for prevention services was in Greater Accra at 88.5% (95%CrI: 86.5-90.2) and highest overall treatment coverage in Upper East at 79.8% (95%CrI: 71.7-86.4). All regions are predicted to achieve the 80% coverage by 2030 with 100% probabilities for composite prevention index; however, the same cannot be applied to composite treatment index as four out of the 10 regions will fail to reach the target, given probabilities less than 80% (Table in S9 Table).

**Figure 3:**
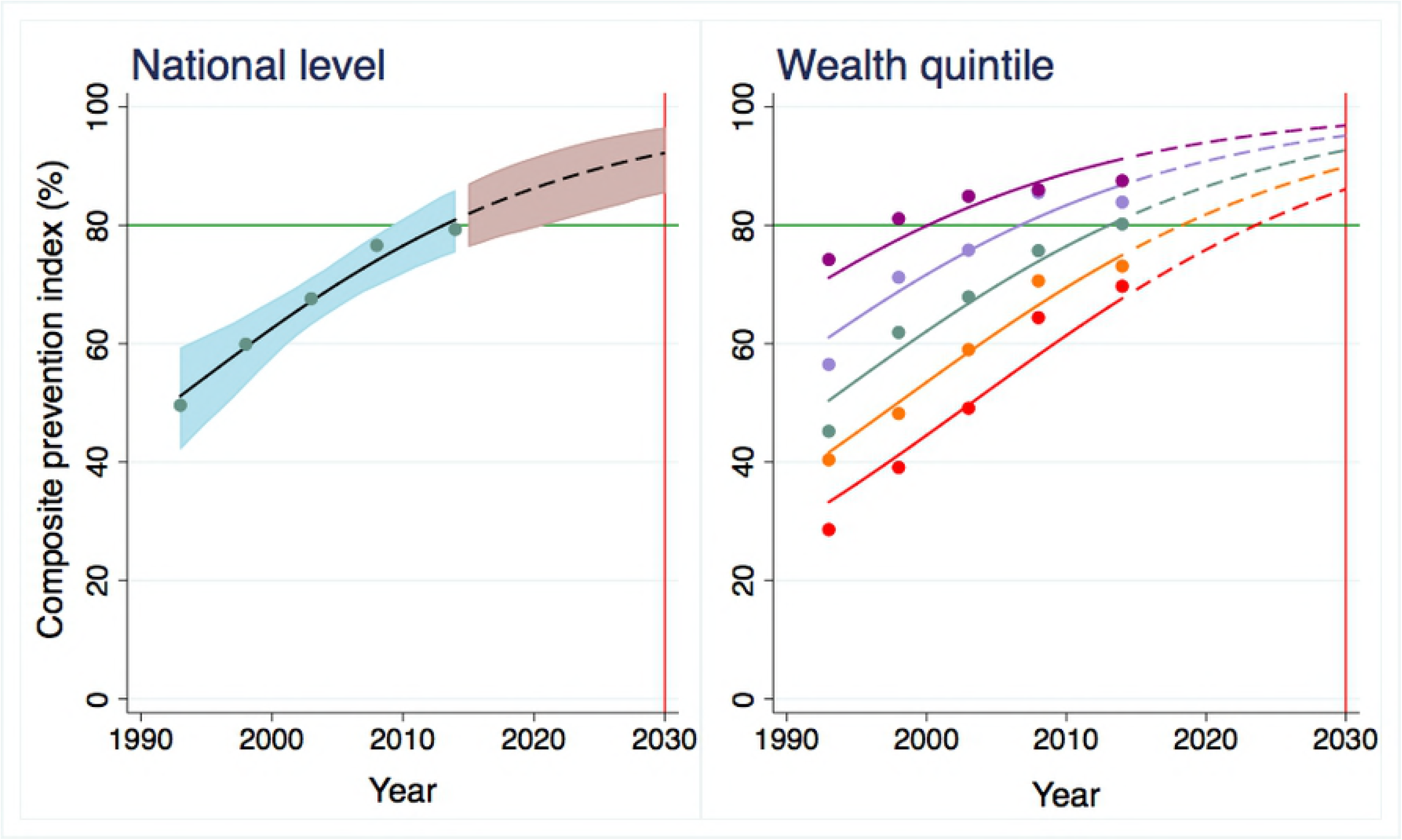
Trends and projections of overall prevention coverage across regions in Ghana, 1993-2030.

**Figure 4:**
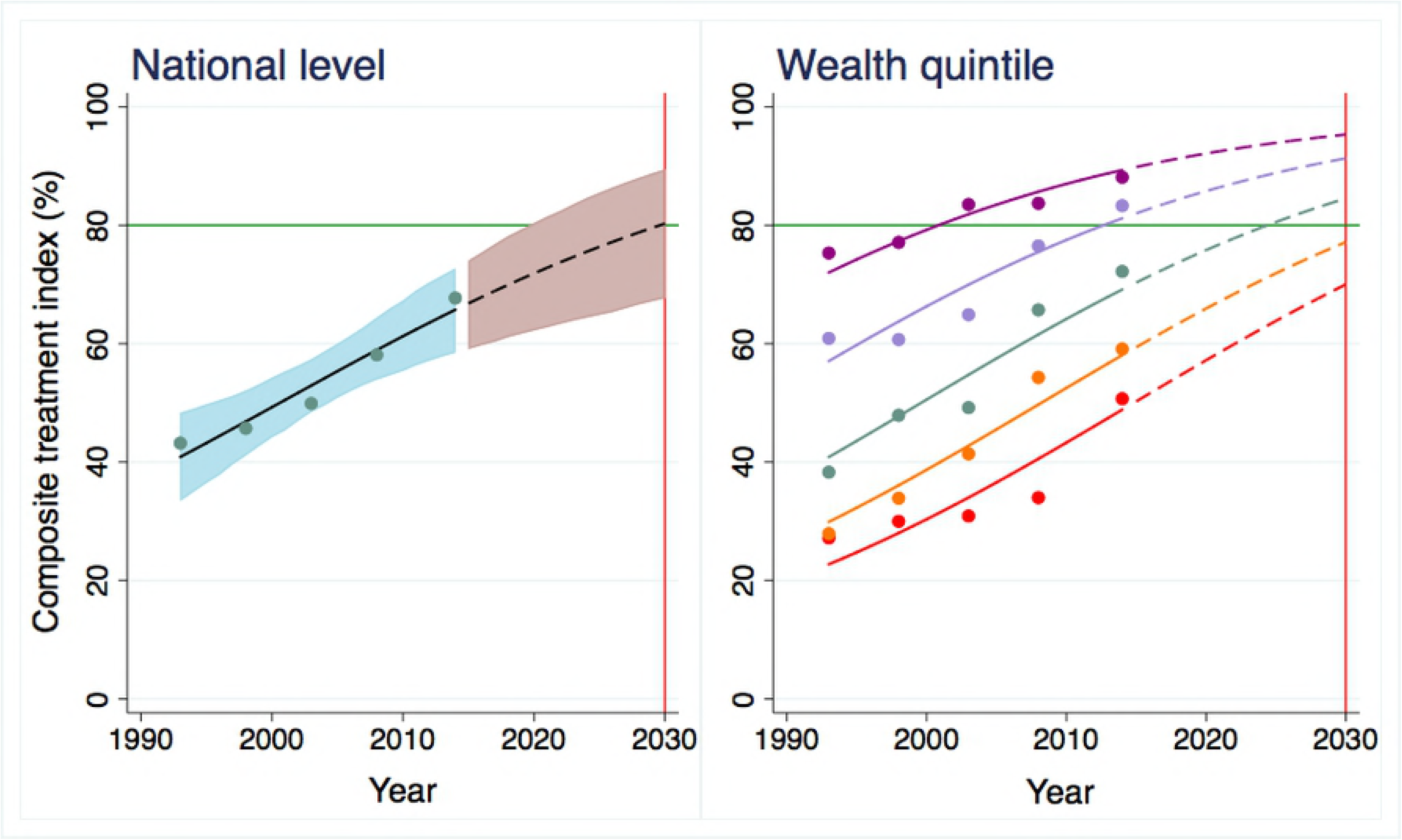
Trends and projections of overall treatment coverage across regions in Ghana, 1993-2030.

### Inequality in service coverage

Table 2 depicts the absolute difference or percentage point difference in the proportion of health service coverage between the extremes of wealth quintiles represented by SII values along with 95% CrI. Overall, absolute inequalities have drastically decreased across most indicators except for use of skilled birth attendance, institutional delivery, and adequate sanitation in which SII only slightly decreased for skilled birth attendance and even increased and will remain so for institutional delivery and adequate sanitation. These three indicators have the biggest inequalities in 2015 and will prevail up to 2030. Adequate sanitation has a SII of 87.7 (95% CrI: 79.4-93.4), followed by institutional delivery at 87.4 (95% CrI: 77.9-93.6) and skilled birth attendance at

**Table 2:** Slope index of inequality (SII) in health service indicators, 1995-2030.

| Indicators | SII (95% CrI) (Q5-Q1)<sup>a</sup> |
| --- | --- |

|  | 1995 | 2005 | 2015 | 2030 |
| --- | --- | --- | --- | --- |
| **Prevention indicators** |  |  |  |  |
| Needs for family planning satisfied | 20.3 (14.6-27.0) | 13.3 (9.7-17.7) | 8.8 (4.1-16.4) | 5.0 (1.0-15.5) |
| At least four antenatal care visits | 52.2 (45.4-59.5) | 40.5 (36.0-45.6) | 29.9 (23.1-38.3) | 17.8 (9.6-30.9) |
| Postnatal care for mothers<sup>c</sup> | — | 75.9 (57.1-89.6) | 54.2 (35.8-70.2) | 25.6 (1.4-77.8) |
| Exclusive breastfeeding | 61.5 (40.5-79.5) | 13.3 (8.7-18.9) | 1.6 (0.4-4.4) | 0.1 (0.0-0.5) |
| Insecticide treated bed nets for children<sup>b</sup> | — | — | — | — |
| Insecticide treated bed nets for mothers<sup>b</sup> | — | — | — | — |
| BCG immunization | 28.0 (19.8-38.4) | 11.2 (8.4-14.7) | 4.1 (2.1-7.3) | 0.9 (0.2-2.6) |
| DPT3 immunization | 40.7 (30.4-52.2) | 14.3 (10.6-18.6) | 4.0 (2.1-6.9) | 0.6 (0.1-1.6) |
| Polio3 immunization | 39.6 (28.6-51.3) | 9.0 (5.8-13.7) | 1.6 (-0.5-4.0) | 0.1 (0.0-0.6) |
| Measles immunization | 38.4 (29.8-48.2) | 15.8 (12.5-19.7) | 5.5 (3.2-8.6) | 1.1 (0.3-2.7) |
| Improved water | 78.7 (71.6-85.8) | 57.4 (39.8-75.0) | 32.5 (16.9-48.1) | 10.5 (4.2-16.7) |
| Adequate sanitation | 71.0 (59.5-80.5) | 80.9 (79.4-93.4) | 87.7 (79.4-93.4) | 93.4 (81.4-98.4) |
| No-use of tobacco<sup>c</sup> | — | 18.2 (4.4-31.9) | 10.5 (2.0-18.9) | 4.4 (0.7-8.2) |
| **Treatment indicators** |  |  |  |  |
| Institutional delivery | 70.9 (58.7-87.1) | 80.7 (73.9-85.9) | 87.4 (77.9-93.6) | 93.0 (79.0-98.5) |
| Skilled birth attendance | 72.5 (62.0-80.9) | 71.4 (64.4-77.5) | 69.9 (56.6-81.1) | 67.0 (40.6-87.7) |
| Oral rehydration therapy | 23.5 (17.8-29.6) | 15.3 (12.6-18.4) | 9.8 (6.4-14.5) | 5.1 (1.9-11.0) |
| Care seeking for pneumonia | 37.0 (26.5-47.8) | 26.9 (21.1-32.8) | 19.0 (11.4-29.0) | 11.3 (3.3-27.9) |

Note: <sup>a</sup>SII= slope index of inequality; Q5 indicates the richest quintile, and Q1 indicates the poorest quintile. <sup>b</sup>unable to obtain accurate trends due to huge variations in SII values from raw data between survey years. <sup>c</sup>No data before year 2000. CrI= credible interval; DPT3= three doses of DPT immunization; Polio3= three doses of polio immunization.

69.9 (95% CrI: 56.6-81.1). These numbers signify that the richest quintile had 69.9 to 87.7 percentage points higher coverage compared to the poor. Although inequalities in indicators related to maternal care such as antenatal and postnatal care have decreased, but persistent inequalities are predicted to exist up to 2030. Similar to absolute inequalities, decreasing trends are also predicted for relative inequalities, represented by RII values, to a lesser degree. By examining current values, it is evident that for most indicators, the richer groups had one to three times more coverage than their poorer counterparts in 2015. Detailed RIIs from 1995 to 2030 is presented in Table in S8 Table.

### Financial hardship

Both incidence of CHE and impoverishment drastically decreased from 1995 to 2030 with an annual rate of reduction at 10.2% (95% CrI: 5.9-14.2) (Table 3). In 1995, 15.0% (95% CrI: 9.6-22.6) of households suffered financial catastrophe as a result of OOP health care payments; however, that proportion will reduce to 0.4% (95%CrI: 0.1-1.3) in 2030. The probability of achieving 100% financial risk protection is estimated to be 96.2%. Proportion of households that experienced impoverishment was 1.7% (95%CrI: 1.1-2.6) in 1995 and is predicted to reduce to

**Table 3:** Trends and projections of the incidence of catastrophic health expenditure and impoverishment in Ghana, 1995-2030.

| Year | Catastrophic health expenditure<sup>a</sup><br>(95% CrI) | Impoverishment<br>(95% CrI) | Financial hardship<sup>b</sup><br>(95% CrI) |
| --- | --- | --- | --- |
| 1995 | 15.0 (9.6-22.6) | 1.7 (1.1-2.6) | 15.5 (10.1-22.9) |
| 2000 | 9.0 (6.4-12.7) | 1.2 (0.8-1.7) | 9.7 (7.0-13.6) |
| 2005 | 5.4 (3.8-7.3) | 0.9 (0.6-1.2) | 5.9 (4.2-8.5) |
| 2010 | 3.2 (2.0-4.9) | 0.6 (0.4-0.9) | 3.6 (2.2-5.6) |
| 2015 | 1.9 (0.9-3.5) | 0.4 (0.2-0.8) | 2.2 (1.1-3.8) |
| 2020 | 1.1 (0.4-2.4) | 0.3 (0.1-0.7) | 1.3 (0.5-2.7) |
| 2030 | 0.4 (0.1-1.3) | 0.2 (0.0-0.5) | 0.5 (0.1-1.4) |
| Annual rate of reduction<sup>c</sup> | -10.2 (-14.2 to -5.9) | -6.7 (-10.7 to -2.7) | -9.7 (-13.6 to -6.0) |
| Probability<sup>d</sup> | 96.2% | 100% | 99.7% |

Note: <sup>a</sup>Catastrophic health expenditure was calculated based on the 25% threshold of nonfood consumption; <sup>b</sup>Financial hardship indicates the incidence of catastrophic health expenditures and/or impoverishment; <sup>c</sup>The annual rate of change for the period 1995-2030; <sup>d</sup>Probability= the probability of meeting the target of 100% financial risk protection by 2030 according to WHO's universal health coverage target; CrI= credible interval.

0.2% (95%CrI: 0.0-0.5) by 2030. Inequality in CHE also decreased from 1995 to 2030 as shown in table 4. In 1995, the poorest quintile was suffering 5.4 percentage points more CHE in comparison to the richest quintile. By 2030, that difference will reduce to 0.1 percentage point.

**Table 4:**
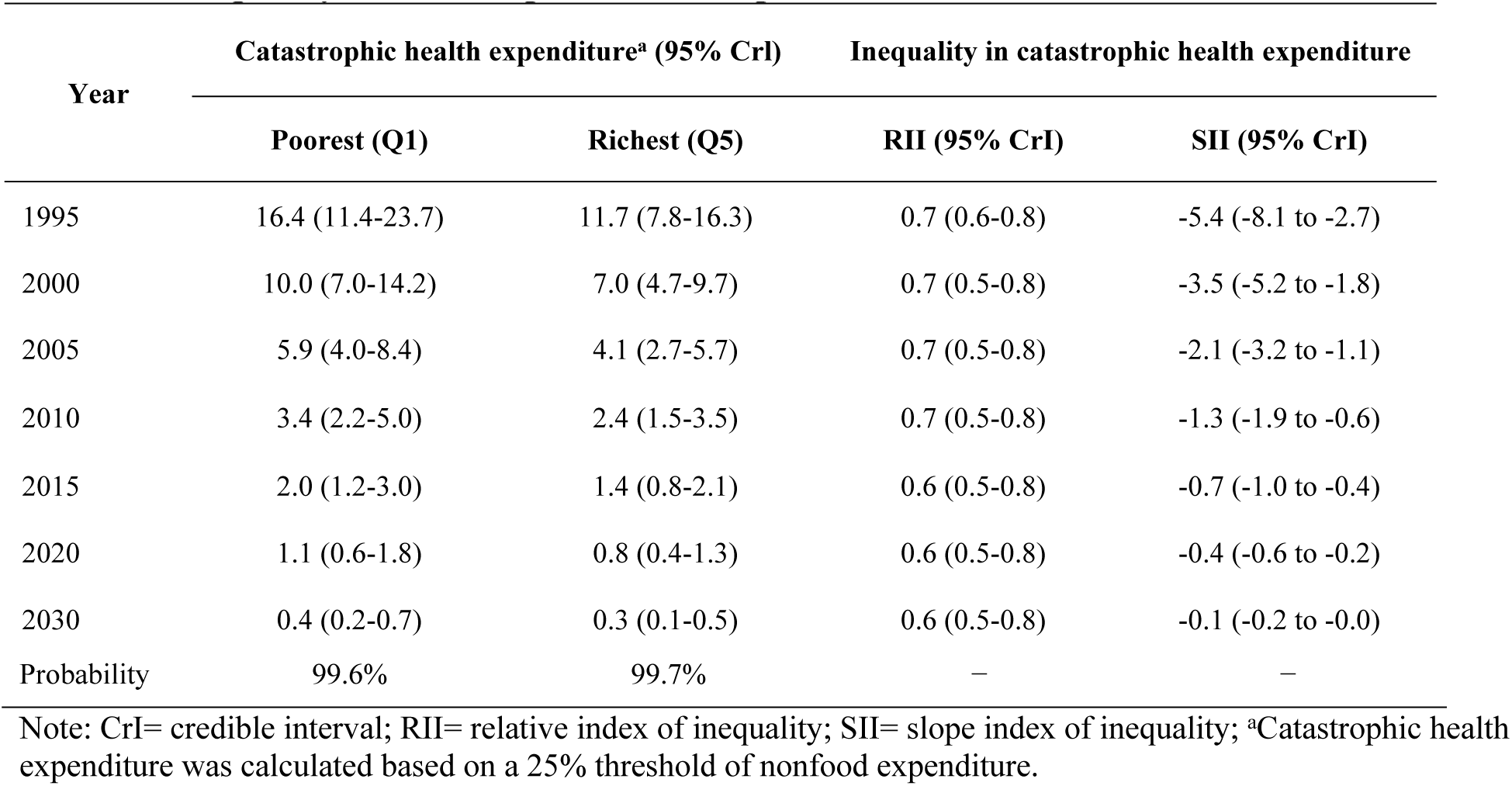
Inequality in catastrophic health expenditure in Ghana, 1995-2030.

At the subnational level as represented in table 5, proportion of households suffering CHE displays a decreasing trend and all regions are estimated to have extremely low incidence by 2030 with remarkably high probabilities of achieving 100% financial risk protection. In 2015, Central region incurred the highest incidence at 2.3% (95% CrI: 1.4-3.5), followed by Volta at 2.1% (95%CrI: 1.3-3.2). At the same time, Brong-Ahafo and Upper East had the highest incidence of impoverishment at 0.8%. Details regarding the incidence of impoverishment for all regions can be found in Table in S10 Table.

**Table 5:** Incidence of catastrophic health expenditure at the subnational level in Ghana, 1995-2030.

| Region | Catastrophic health expenditure<sup>a</sup> (95% CrI) |  |  |  | Probability<sup>b</sup> |
| --- | --- | --- | --- | --- | --- |
|  | 1995 | 2005 | 2015 | 2030 |  |
| Ashanti | 16.1 (10.5-22.5) | 5.9 (3.8-8.6) | 2.0 (1.2-3.1) | 0.4 (0.2-0.7) | 99.5% |
| Brong-Ahafo | 14.6 (9.5-20.3) | 5.3 (3.4-7.6) | 1.8 (1.1-2.8) | 0.4 (0.2-0.6) | 100% |
| Central | 17.9 (11.9-24.7) | 6.7 (4.3-9.5) | 2.3 (1.4-3.5) | 0.5 (0.2-0.8) | 99.7% |
| Eastern | 15.0 (10.2-21.2) | 5.5 (3.6-7.9) | 1.9 (1.1-2.9) | 0.4 (0.2-0.7) | 99.9% |
| Greater Accra | 9.5 (6.2-13.7) | 3.3 (2.2-4.9) | 1.1 (0.7-1.7) | 0.2 (0.1-0.4) | 100% |
| Northern | 13.0 (8.6-18.5) | 4.7 (3.0-6.8) | 1.6 (1.0-2.5) | 0.3 (0.2-0.6) | 100% |
| Upper East | 9.1 (5.9-13.4) | 3.2 (2.1-4.7) | 1.1 (0.6-1.7) | 0.2 (0.1-0.4) | 100% |
| Upper West | 8.4 (5.3-12.5) | 2.9 (1.8-4.4) | 1.0 (0.6-1.5) | 0.2 (0.1-0.3) | 100% |
| Volta | 16.3 (11.2-22.9) | 6.0 (4.0-8.7) | 2.1 (1.3-3.2) | 0.4 (0.2-0.7) | 99.6% |
| Western | 13.8 (9.1-19.9) | 5.0 (3.2-7.2) | 1.7 (1.0-2.7) | 0.3 (0.2-0.6) | 99.9% |

Note: <sup>a</sup>Catastrophic health expenditure was calculated based on 25% threshold of nonfood expenditure; <sup>b</sup>The probability of achieving 100% financial risk protection; CrI: credible interval

## Discussion

This paper provides a comprehensive picture of Ghana’s progress towards UHC, as it extensively examined both health service coverage and financial risk protection components at the national, subnational, and wealth quintile levels. There have been tremendous improvements in increasing access to various health services with upward trends in coverage accompanied by decreasing trends in the proportion of households suffering CHE. Even though Ghana implemented the NHIS in 2003 as an attempt to remove barriers to cost and increase health service utilization especially among the poor [16], inequalities still exist for both UHC components throughout the nation. Similarly, subnational disparities were apparent as well with some regions faring better than others.

Findings from the analyses indicate that most of the maternal health indicators such as more than four antenatal care visits, postnatal care for mothers, institutional delivery, and use of skilled birth attendance have already reached the 80% target in 2015 at the national level, most likely due to the implementation of NHIS as this insurance allows pregnant women to be exempt from premium with maternity care coverage [17]. In addition, following enrollment into NHIS, the Maternal Health Program, granted to women upon confirmation of pregnancy, completely eliminates OOP payments for six antenatal care visits, delivery care, two postnatal care visits, and infant care up to three months of age [17]. This could be a reason why postnatal care for mothers increased by eight-fold in the span of 10 years from 2005 to 2015 and high coverage of childhood vaccinations. Countries such as Rwanda and Indonesia have also proved that pregnant women who are enrolled into insurance are more likely to utilize various maternal care services such as prenatal care, institutional delivery, skilled birth attendants, postnatal care, and seek vaccinations for their children [16-18]. Therefore, continuous efforts must be made to encourage pregnant women to enroll into NHIS and to utilize maternal care services.

At the national level, family planning demand satisfied showed the least likelihood of reaching its target and the reason could be attributed to the fact that this service is not covered under NHIS but implemented through a government vertical program [19] and this pattern is consistent with a previous study [20]. The low uptake of family planning can be explained by many facets such as misconception, stigma, lack of knowledge, religious abhorrence, spousal disapproval, and inaccessibility [21]. To increase the uptake of family planning, national efforts to include it as a component of the maternal health care package under the NHIS, and education and involvement of male partners should be considered. Community health workers (CHWs) can greatly improve access to contraceptives and provide education on their correct use [22].

Similarly, care seeking for the treatment of pneumonia also had a low coverage and unlikely to reach the target because a past study conducted in rural Ghana found significant knowledge deficit among residents regarding pneumonia [23]. In that study, only one-third of the studied population ever heard of the disease name and among those, only half sought treatment for their children [23]. It is imperative to increase people’s knowledge of childhood illnesses and perhaps the Ministry of Health and Ghana Health Service need to strengthen the health promotion unit to provide structured and targeted community educational programs. Enrolling into NHIS can potentially improve treatment-seeking behavior since parents who are enrolled are more likely to seek curative and preventive care for their children [24].

We also found that utilization of institutional delivery and access to skill birth attendance were among the top three indicators with the biggest absolute and relative inequalities in 2015. Several studies have found that wealthier and more educated women are more likely to enroll into NHIS and seek maternal care services [17-19]. Many women from poor households lack the basic knowledge about insurance system such as not being aware of the fact that their children and maternal health services are exempt from insurance premium; some simply did not know how to register themselves or their children [17]. This further highlight the need for more community based education on the NHIS targeting especially women from poorer households to enroll.

Community health workers (CHWs) have proven to be effective in improving health care access and overall health status [25,26]; therefore, they can also be used to educate and encourage enrolment of caregivers and mothers onto the NHIS. These health workers can act as channels in reaching out to the poor women and educate them on the benefits package for maternal health under the NHIS and encourage them to enroll.

The study found wealth quintile based differences in coverage among most health service indicators, which indicated that enrollment into insurance maybe concentrated among the rich. This observation is consistent with findings in previous studies [18,27,28]. Furthermore, a study done in rural Ghana found that enrollment into NHIS is 2.5 times more among the rich than the poor [29]. This deviates from NHIS’ original equity goal, which was to increase affordability and utilization of health services especially among the poor and vulnerable population [19]. If the poor are not insured, there is an increased likelihood of illnesses resulting in worse health status and further impoverishment. This will not only widen inequality gap but also drifts away from UHC goals. In addition to quintile-based differences, inequality also prevailed at the subnational level with Northern region lagging significantly behind all other regions in prevention and treatment coverage. This could be explained by the fact that this region occupies about a third of the land-size of Ghana, and has the lowest doctor-to-population and nurse-to-population ratios, making health services more difficult to obtain [17]. Furthermore, this region is the biggest contributor to poverty in Ghana [30]. Overall, regional specific strategies (administrative and social policy) will be needed. Some policy measures such as budgeting of CHWs, and promoting current partnership with One Million CHW Campaign [25] will be keys. Wealth, education, employment, and location play critical roles in health service coverage, thus national and sub-national efforts are needed to increase all aspects in order to improve coverage [31].

Financial catastrophe decreased by almost eight-fold from 1995 to 2015, proving that NHIS have aided in reducing OOP health payments across the nation. During the same period, impoverishment witnessed a four-fold decrease. Although inequality gaps have narrowed as difference in the incidence of financial catastrophe between rich and poor groups reduced, the poor is still suffering more CHE as a result of OOP payment. This finding is in congruence with previous studies from other Sub-Saharan African countries [32-34]. Moreover, benefits received from health care services are also unequal among those paying for it. In Ghana, benefits from the public sector and private services tend to be pro-rich despite more health care needs by the poor [33]. A study found that 23% of those indicated having poor health in the poorest quintile only received less than 13% of total health care benefits compared to 16% of those in the richest quintile obtaining over 24% benefits [6]. At a threshold of 25% non-food expenditure, incidence of CHE was 1.8% in 2015 and 0.4% of households were pushed into poverty. These results were still worse compared to earlier studies done in South Africa and Tanzania in which impoverishment were only 0.045% and 0.37%, respectively in 2008 [33]. On a positive note, Ghana’s progress was shown to be better than Rwanda and Nigeria [34-36].

From these results, it is evident that NHIS has contributed to Ghana moving one step closer to realizing UHC with great improvement seen in both components of the UHC. However, persistent inequalities between rich and poor were apparent at all levels in health service coverage and financial risk protection. Insurance premiums and registration fees remain as obstacles to enrolment for the poor, many are simply left unattended without any form of protection [27]. For people in the poorest quintile, premium imposes a heavy burden as it takes up 11.4% of their non-food expenditure while it only takes up 5.9% for general households [28]. A recent study done in Ghana revealed that as share of OOP payment for healthcare in total household expenditure increases, impoverishment deepens [37]. Reducing OOP payment for healthcare, especially for the poor, is essential as it is a determining factor of impoverishment. UHC policy has proven to greatly contributes to the reduction of catastrophic payments [38].

Regardless of NHIS’ shortcomings, it is still considered an accomplishment for Ghana, making the country one of the few in Sub-Saharan Africa to implement a nationwide insurance scheme. Previous studies conducted in Ghana regarding the protective effect of NHIS revealed that incidence and intensity of CHE were greatly reduced in insured individuals especially for the poor [29,39]. It proved to have protective mechanism against financial shocks with a 67% reduction in OOP and reduces the likelihood of foregoing other subsistent needs for health care [29]. It is recommended that Ghana to continue with its current progress by further enrolling all citizens into the scheme. There is a need to explore other funding options to ensure sustainability of NHIS as well as to further lower premium for the poor and vulnerable population. Thailand sets a great example in establishing equity through its insurance scheme by subsidizing tax for the poor instead of premium contribution [40]. It is believed that free health care for all is achievable and affordable in Ghana via cost savings, progressive taxation, and high quality transparent aid [41].

## Strengths and limitations

One of the major strengths of this study is that it comprehensively examined both health service coverage and financial risk protection. It also provided trends and future projections for chosen indicators. Furthermore, subnational analysis yielded further detailed assessment. Few accompanied limitations include missing information prior to 2000 for some of the indicators, the inability to make projections for NCD and HIV related indicators, and the inability to capture information for those that are too poor to utilize healthcare.

## Conclusion

Ghana with its strenuous efforts in making health care more equitable and affordable has made tremendous improvements in health service coverage along with reducing out-of-pocket (OOP) payment owing to the enforcement of NHIS. The establishment of NHIS was also an attempt to achieve MDGs, especially in the area of maternal and child health in which significant improvements were observed. However, apparent inequalities were evident at the national and subnational level since the poor were suffering more catastrophic health expenditure (CHE) and had less access to health services. These inequalities were observed in many studies conducted in Ghana thus far. Policy makers need to have stronger commitments in achieving equity as many health care interventions, aimed at the poor, do not reach them. Several options to be considered are equitably distribute funds to regions according to needs, reduce copayment by exempting or heavily subsiding premium for the poor, and enhance Community-based Health Planning and Service programme to improve access to hard-to-reach population. Ghana serves as an exemplary example for other Sub-Saharan African countries in implementing a health insurance at the national level. Its achievement in improving health care utilization for its citizens and reducing financial burdens is praiseworthy. It is recommended that Ghana to continue with its current progress by further enrolling all its citizens into the scheme. By doing so, UHC will no longer be a distant dream.

## Supporting information

**S1 Table: Health service indicators**

**S2 Table: Deviance information criteria for health service indicators**

**S3 Table: Quintile-specific coverage of reproductive, maternal, and child health services in Ghana, 1995-2030**

**S4 Table: Quintile-specific vaccination coverage in Ghana, 1995-2030**

**S5 Table: Quintile specific coverage of disease prevention and environmental health indicators in Ghana, 1995-2030**

**S6 Table: Quintile specific coverage of treatment services for delivery care and childhood illnesses in Ghana, 1995-2030**

**S7 Table: Quintile-specific coverage of composite indices for health services in Ghana, 1995-2030**

**S8 Table: Relative index of inequalities (RII) in health service indicators, 1995-2030**

**S9 Table: Overall prevention and treatment service coverage at the subnational level in Ghana, 1995-2030**

**S10 Table: Impoverishment at the subnational level in Ghana, 1995-2030 S11 Table: Survey characteristics**

**S1 Figure: Trends and projections of composite coverage index in Ghana, 1993-2030**

**S2 Figure: Trends and projections of the coverage of insecticide treated bed nets for children under five and pregnant women in Ghana, 2003-2030**

**S3 Figure: Trends and projections of the coverage of antenatal and postnatal care for women in Ghana, 1993-2030**

**S4 Figure: Trends and projections of the coverage of family planning needs satisfied and exclusive breastfeeding in Ghana, 1993-2030**

**S5 Figure: Trends and projections of the coverage of delivery care services in Ghana, 1993-2030**

**S6 Figure: Trends and projections of the coverage of measles and polio vaccination in Ghana, 1993-2030**

**S7 Figure: Trends and projections of the coverage of BCG and DPT vaccination in Ghana, 1993-2030**

**S8 Figure: Trends and projections of the coverage of treatment services for childhood illnesses in Ghana, 1993-2030**

**S9 Figure: Trends and projections of the coverage of environmental health indicators in Ghana, 1993-2030**

**S10 Figure: Trends and projections of non-tobacco users in Ghana, 2003-2030**

## Acknowledgements

The author thank the Department of Global Health Policy at The University of Tokyo, School of Public Health at the University of Ghana, professors, assistant professor, and fellow lab mate for their assistance. The author also thanks the Ghana Statistical Service for surveys and data.

